# Increasing the acquisition speed in oblique plane microscopy via Aliasing

**DOI:** 10.1101/2024.12.25.630337

**Authors:** Conor Mcfadden, James Manton, Reto Fiolka

## Abstract

Oblique plane microcopy (OPM), a variant of light-sheet fluorescence microscopy (LSFM), enables rapid volumetric imaging without mechanically scanning the sample or an objective. In an OPM, the sample space is mapped to a distortion free image space via remote focusing, and the oblique light-sheet plane is mapped via a tilted tertiary imaging system onto a camera. As a result, the 3D point-spread function and optical transfer function are tilted to the optical axis of the tertiary imaging system. To satisfy Nyquist sampling, small scanning steps are required to encompass the tilted 3D OTF, slowing down acquisition and increasing sample exposure. Here we show that a judicious amount of under-sampling can lead to a form of aliasing in OPM that can be recovered without a loss of spatial resolution or introducing artifacts. The resulting speed gain depends on the optical parameters of the system and can reach 2-4-fold in our demonstrations. We leverage this speed gain for rapid subcellular 3D imaging of mitochondrial dynamics.

## 1. Introduction

Light-sheet fluorescence microscopy (LSFM), owing to its low sample irradiance and rapid volumetric imaging capabilities has found numerous applications in biological and biomedical research[1]. By illuminating the sample with a sheet of light, intrinsic optical sectioning is achieved, which limits unnecessary sample irradiation outside the focal plane and allows rapid image acquisition. In a typical light-sheet microscope, illumination and detection are implemented on separate objective lenses, typically oriented orthogonally to each other. 3D image acquisition is either performed by scanning the sample through the light-sheet, or by optically scanning the light-sheet and mechanically translating the detection objective [1].

Oblique plane microscopy [2] (and related techniques such as Swept confocally-aligned planar excitation, SCAPE [3]) is a type of LSFM where both illumination and detection are performed by a single primary lens. This improves sample accessibility, and in recent implementations of OPM using rapid optical scanning technology [4, 5], much higher volumetric acquisition speed can be obtained than using traditional LSFM architectures.

In a traditional wide-field microscope, the principal axes (x, y, z) of the point spread function (PSF) are aligned with the principal axes (s_x_, s_y_, s_z_) of sampling (i.e. along the optical axis and in the axes of the pixel array on the camera). In an OPM, because the light sheet is launched from the primary objective at an oblique angle, the image plane is also tilted by an oblique angle. As such, for an OPM, both s_x_ and s_z_ have components in the x and z directions. As a result, lateral and axial resolution are coupled and relatively fine step sizes are needed to acquire 3D data. This in turn slows down 3D imaging and burdens the sample with enhanced radiation exposure, compared to an imaging system where the OTF is aligned with the optical axis.

Here we show that some degree of under-sampling in OPM leads to a form of frequency aliasing that can be recovered without a loss of spatial resolution while minimizing artifacts. This is because the tilted OTF leaves some voids in 3D Fourier space which can accommodate aliased information without overlap.

The amount of tolerable under sampling, and resulting gain in imaging speed, depends on the properties of the imaging system. We demonstrate two-fold gains for a high resolution OPM and 4-fold speed gains for a mesoscopic system. We leverage the method to achieve rapid volumetric imaging of mitochondrial dynamics at two-fold higher speeds than using critical Nyquist sampling.

### 2.1 Theory

**Figure 1A** shows schematically the principle of OPM. A light-sheet (blue) emerges at an oblique angle from the primary objective (O1). It excites two emitters (green dots), whose fluorescence emission is partially captured by the primary objective (green cones). A remote focusing system [6] maps the image of the emitter via a matched secondary objective (O2) into a remote space. Due to the limited collection angle, the 3D image of an emitter, i.e. its point spread function, is elongated along the optical axis (z). The 3D PSF is invariant (within a finite axial range [6]) within the remote focus volume, so both emitter images share the same PSF, even though the originate from different depths. A tertiary imaging system (Objective O3), tilted to the same angle as the lights-sheet, maps the fluorescence emission onto a detector [2]. By probing the remote space at an angle, the point spread function appears tilted in the tertiary reference frame (X’-Y’-Z’, see also **Figure 1B**). We ignore here effects of only capturing a subset of the light-cone emitted by O2 with O3, which may introduce an additional source of PSF tilt.

**Fig. 1.**
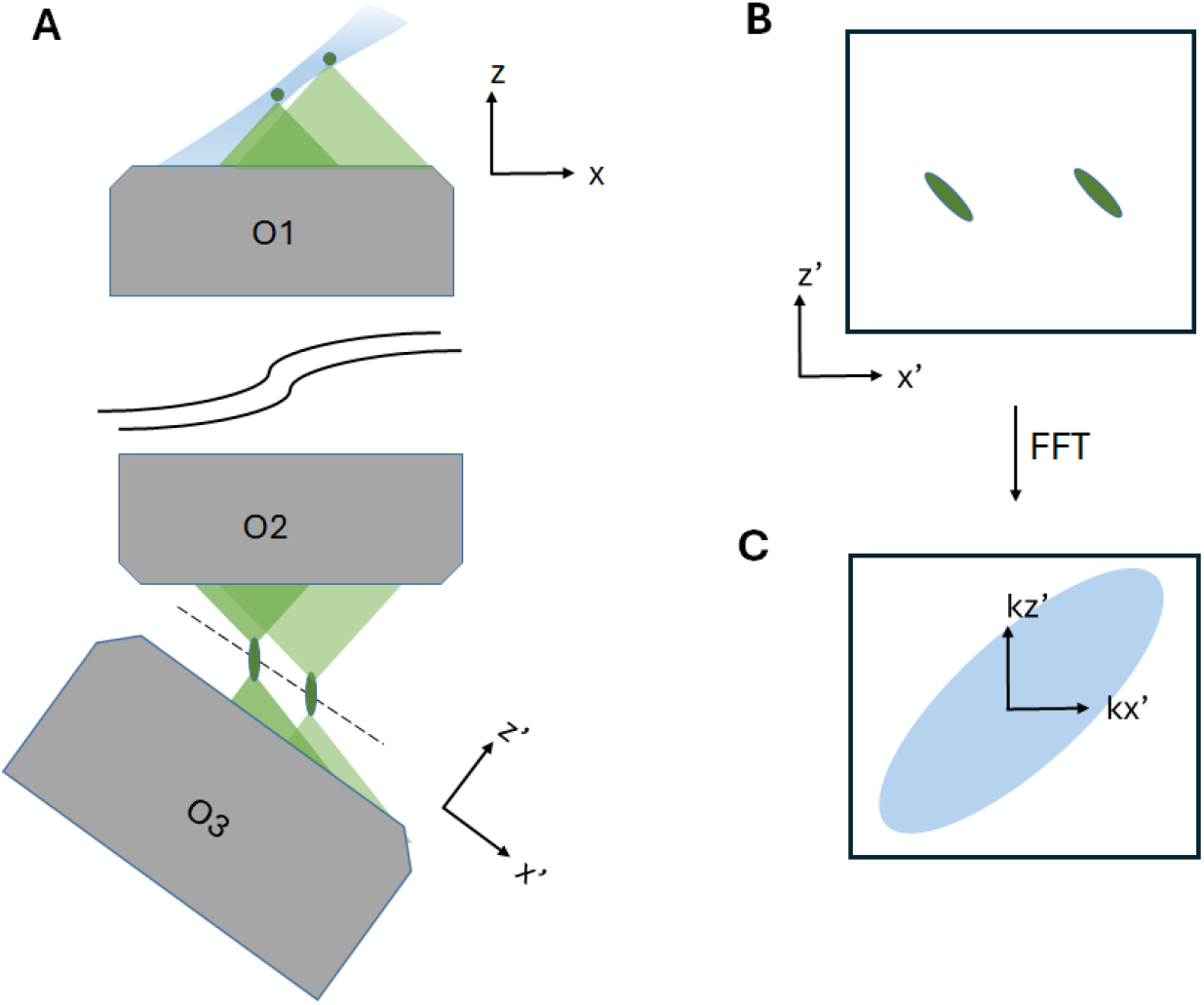
Schematic illustration of an OPM, and its imaging properties. **A** Simplified optical train in OPM: A tilted light-sheet (blue) emerges from the primary objective (O1). The fluorescence light from two emitters (green dots) is relayed via the secondary objective (O2) to the tilted tertiary objective (O3). **B** Volumetric image of the two emitters, as acquired in the reference frame of the tertiary objective (Z’-X’). **C** Fourier transform of the volumetric dataset of the emitters shown in **B**. The optical transfer function is represented in light-blue.

A Fourier transform of the 3D PSF yields the optical transfer function, which is correspondingly also tilted. For simplicity, we assume that the OTF has an ellipsoidal shape, with the magnitude of its major axes scaling proportionally to the resolving power. Typically, the lateral resolution (i.e. X and Y direction) is 3-4 fold better than in the third dimension (Z direction), and hence the OTF has a similar aspect ratio. We note that since the overall PSF is the product of the light-sheet intensity distribution and the detection PSF, the OPM PSF is slightly skewed due to the light-sheet tilt (i.e. the light-sheet propagation axis and the Z axis are not orthogonal to each other, as in a traditional LSFM).

**Figure 2A** illustrates critical Nyquist sampling requirement for the lateral, p_x’_ and axial, p_z’_ sampling steps (i.e. voxel sizes). For the sake of argument, we assume a 45-degree tilt of the OTF, and an aspect ratio of K_x_/K_z_=4, with K_x_ and K_z_ being the cut-off frequencies of the OTF in the lateral and axial direction, respectively. The Fourier transform of the pixel grid with p_x’_ and p_z’_ voxel size in real space corresponds to a grid with 1/p_x’_ and 1/p_z’_ spacing in reciprocal space. At every grid point, a copy of the OTF is located. In order to avoid overlap of the OTF copies, we find that p_x_’=p_z_’, and that their magnitude needs to exceed Kx/sqrt(2). Then the borders of the numerical domain of a discrete Fourier transform (black box in **Figure 2A**) encompass the central copy of the OTF, and no neighboring copy of an OTF overlaps with that domain.

**Fig. 2.**
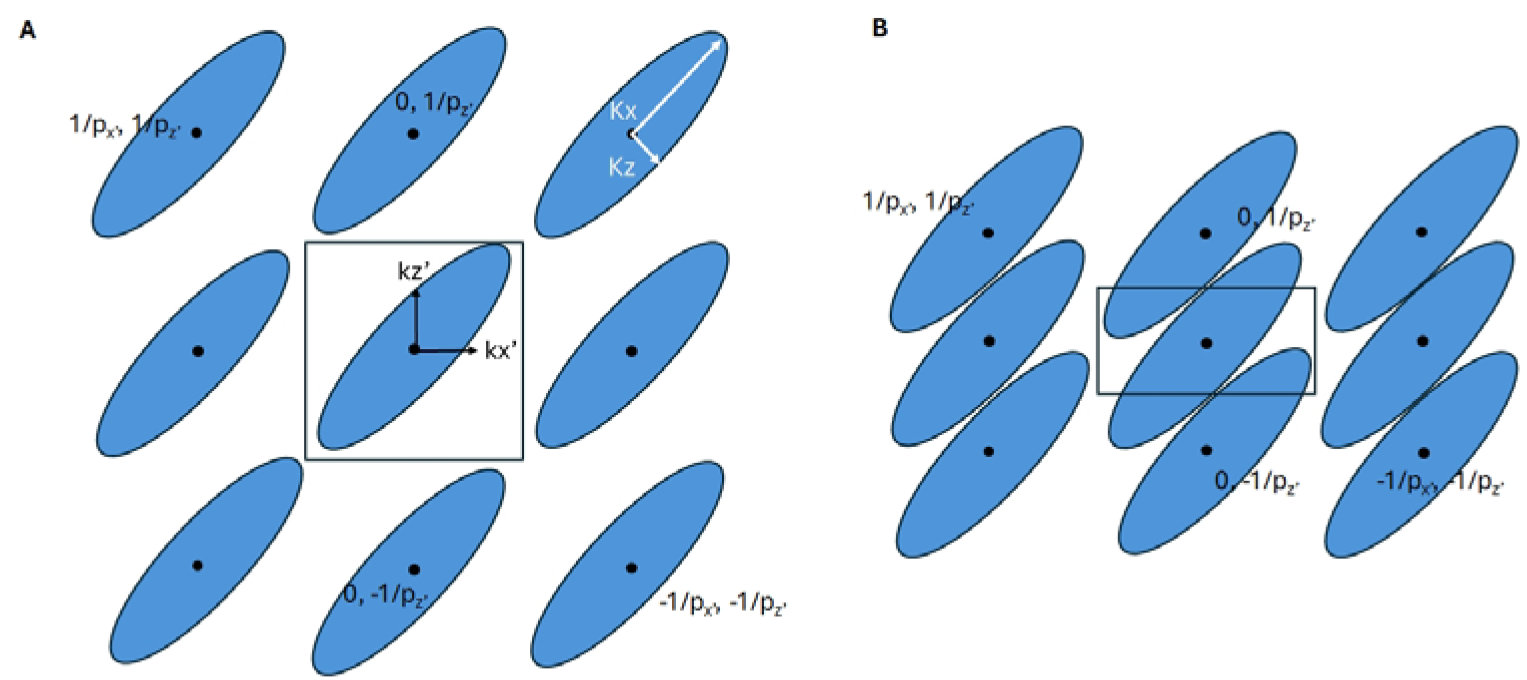
Sampling of the tilted OPM OTF. **A** Critical Nyquist sampling of the OPM OTF (blue ellipses). **B** Axial undersampling by a factor of 2. p_x_, p_z_: lateral and axial pixel size. Kx,Kz: cut-off frequencies of the OTF.

If we double the axial step size, we expect to violate the Nyquist condition. Indeed, in **Figure 2B**, one can see that the grid in reciprocal space has contracted by a factor of 2 in the k_z’_ direction. Because of the under sampling, not all frequency components of the central copy of the OTF are contained within in the numerical domain of a discrete Fourier transform. Furthermore, parts of two OTF copies, located at 0,1/p_z_’ and 0,-1/p_z_’, have leaked into the numerical domain.

Importantly, there is no overlap between OTF copies themselves in the scenario shown in **Figure 2B**, but there is a misplacement of Fourier components. We reasoned if we could properly re-arrange the Fourier components, a proper full support of the OTF could be obtained, which is inspired by how structured illumination microscopy (SIM) reassigns misplaced Fourier components [7]. However, in contrast to SIM, there are no overlapping Fourier components, thus no prior unmixing needs to be performed.

Our proposed reconstruction scheme is shown in **Figure 3**. In **Figure 3A**, the actual observed information in a 2x under-sampled dataset in Fourier space is shown. We reasoned that if we concatenated three copies of the dataset in the k_z_ direction in reciprocal space, the re-assignment of the Fourier components for the central copy of the OTF would be achieved (see also **Figure 3B**). Obviously, there are still additional copies of the OTF that would cause severe aliasing in real space. However, since those components are separated, they can be removed with a masking step, as illustrated in **Figure 3C**.

**Fig. 3.**
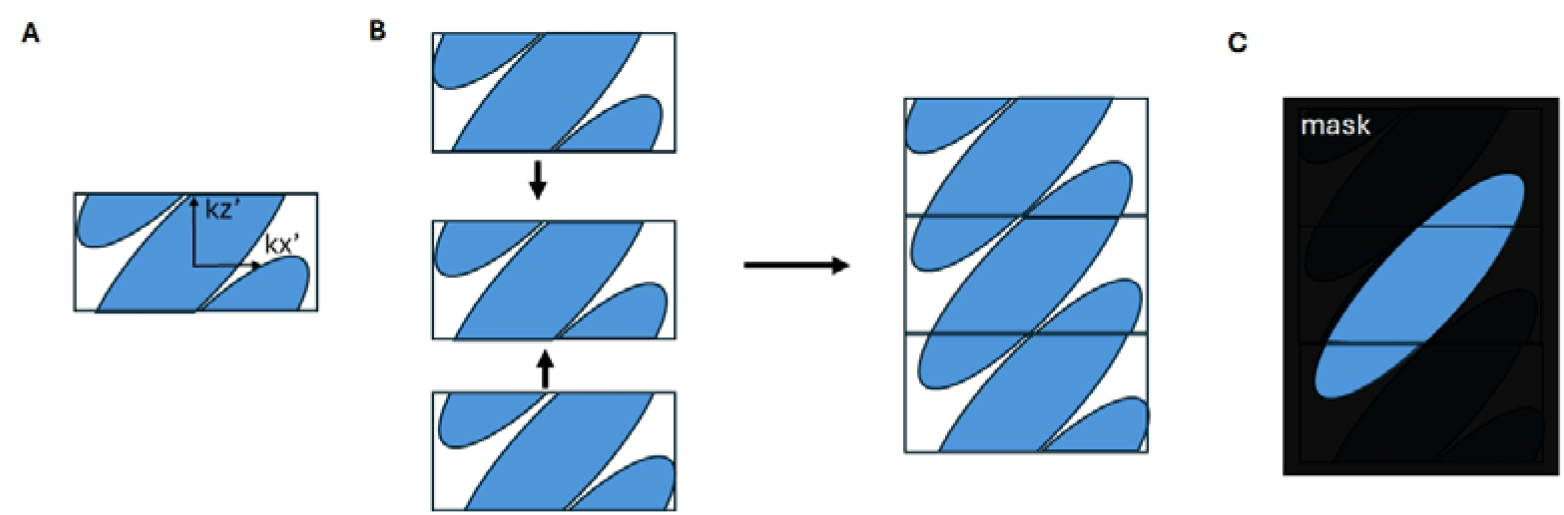
Reconstruction of OPM OTF after under-sampling. **A** Fourier transform of an axially under sampled OTF. **B** Reconstruction of the OTF by concatenating three copies of the OTF along the k_z_ direction. **C** Masking of the central OTF.

The amount of under sampling that can be applied in our technique depends on the tilt angle and aspect ratio (i.e. K_x_/K_z_) of the OTF (see also **Supplementary Figure S1**). In this manuscript, we empirically found the tolerable under-sampling factors for the OPM systems used: a heuristic criterion was the axial resolution, which starts to deteriorate when the under-sampling factor becomes critical as then the aliased copies of the OTF begin to overlap (**Supplementary Figure S2**). Importantly, one can in principle run higher under sampling factors, at the cost of a degradation of axial resolution (**Supplementary Figure S2 and S5)**.

## 2. Results

### 2.1 Fluorescent nanospheres

We have acquired 3D datasets with two OPM systems to demonstrate our method. In **Figure 4A**, the OTF of a high resolution OPM system is shown (40X NA 1.25 Silicone oil objective primary lens, 45-degree light-sheet tilt, unpublished system), where we adjusted the step size to critical Nyquist sampling. In **Figure 4B**, an OTF with 2X under sampling is shown. The red arrows point at two copies of the OTF that have been mixed in via aliasing. These two copies come close to the central body of the OTF, but do not overlap with it yet. **Figure 4C** shows a three-fold repeat of the under sampled OTF shown in **Figure 4B**. As predicted in **Figure 3**, a full central OTF is reconstructed, with additional partial copies flanking it. For reconstruction, we only kept the Fourier components within the two red lines shown in **Figure 4C** and set the rest to zero outside of that band. This represents a top hat (sinc in real space) filter, which may cause some ringing in real space. Alternative windowing methods could be applied, but for this first proof of principle, we employed top hat filtering only. The full workflow is also shown in **Supplementary Figure S3**.

**Figure 4.**
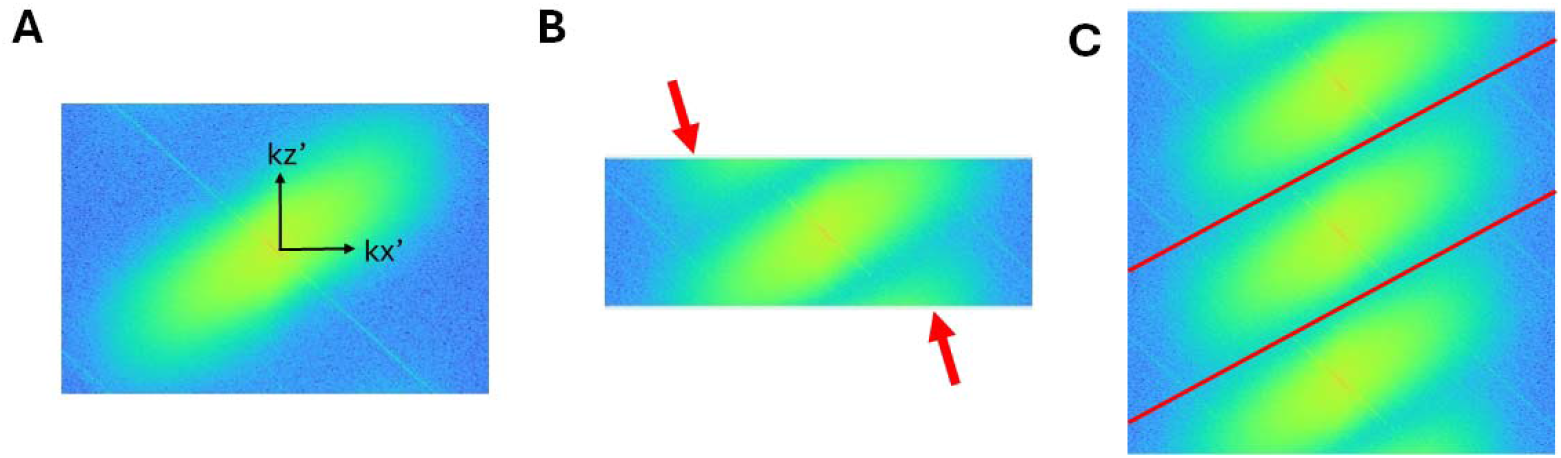
Experimental OPM OTFs, shown as a cross-section and represented as the logarithm of the power spectrum. **A** Critically sampled OTF of a high resolution OPM system. **B** Two-fold under-sampled OTF. The arrows point at Aliasing of OTF components. **C** Threefold repeat of the under sampled OPM OTF. The red lines outline the masking region: All Fourier components outside the band encompassed by the red lines were set to zero.

In **Figure 5**, we compare the real space results for fluorescent nanospheres, as imaged with a high resolution and a mesoscopic (termed meso) OPM. For the high resolution OPM, we imaged 100nm fluorescence nanospheres on a coverslip. The data was de-skewed and rotated using the PetaKit5D software [8], and is presented in an X-Y coordinate frame, projected along the X direction. **Figure 5A** shows the ground truth dataset acquired with critical Nyquist sampling (as shown in **Figure 4A**). **Figure 5B** shows the same beads, but with 2X under-sampling. Typically, undersampled datasets in OPM are interpolated at the rotation step to achieve the proper z-step size, but this leaves pronounced aliasing artifacts in place (red arrows). **Figure 5C** shows our reconstruction from the under-sampled data, in which such artifacts are noticeably absent.

**Figure 5.**
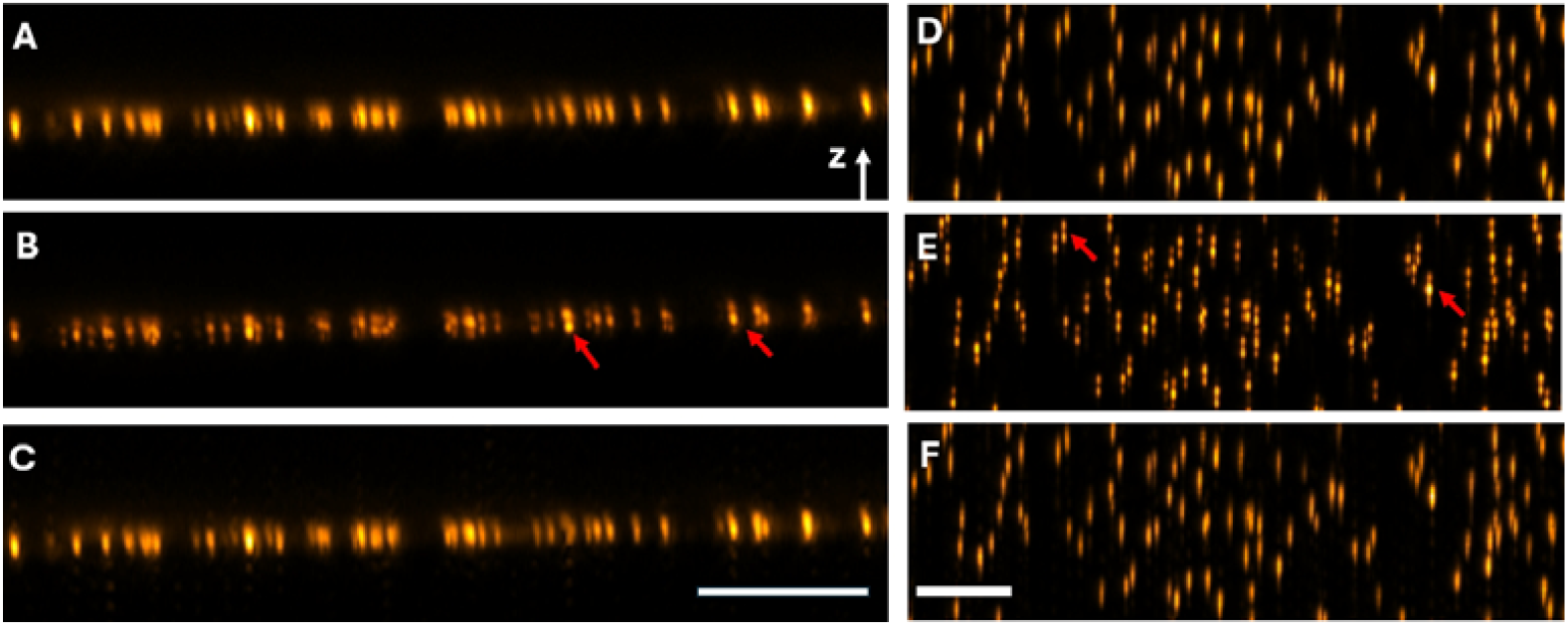
Point Spread Functions for OPM. **A** Critically sampled PSFs for a high resolution OPM. **B** The same region, but acquired with 2x under-sampling. **C** Reconstruction from the 2x under-sampled data. **D** Critically sampled PSFs for a mesoscopic OPM. **E** The same region, but acquired with 4x under-sampling. **F** Reconstruction from the 4x under-sampled data. The red arrows point at artifacts in the under-sampled PSF, such as bifurcation, or concentration into a single spot. Scale bars: **C**: 10 microns; **F** 50 microns.

For the meso OPM using a previously published system [6], we imaged 500nm fluorescent nanospheres in Agarose. We empirically found that 4X under-sampling is possible (**Supplementary Figure S2** and **S3**). **Figure 5D** shows a ground truth data set acquired with critical Nyquist sampling, whereas **Figure 5E** shows the same area, but for 4X under-sampling. Red arrows point at aliasing artifacts, such as bifurcation and concentration into a single spot. **Figure 5F** shows our reconstruction of the 4X under sampled dataset, where the artifacts seen in **Figure 5E** have been removed.

### 2.2 Rapid imaging of mitochondrial dynamics

Next we applied our accelerated acquisition scheme to rapid imaging of subcellular dynamics using the high resolution OPM system. We imaged U2OS cells labeled with OMP25-GPF, an outer membrane marker for mitochondria, with 2X under-sampling (same settings as in **Figure 5A-C**) yielding a volume rate of ∼2Hz. **Figure 6A** shows a cross-sectional view after conventional OPM post-processing (de-skewing, rotation into X-Y-Z reference frame, and interpolation of the Z-axis). The red arrows point at similar aliasing artifacts as seen in **Figure 5B**, such as bifurcation or concentration of the PSF. **Figure 6B** shows the same cross-section, but reconstructed using our method, which is free of those artifacts. We also acquired a separate stack at full sampling for a ground truth comparison (**Supplementary Figure S4**), which was also free of artifacts. Digital down-sampling (removing slices from the stack before further processing) produced artifacts of similar appearance to the once shown in **Figure 6A**.

**Figure 6.**
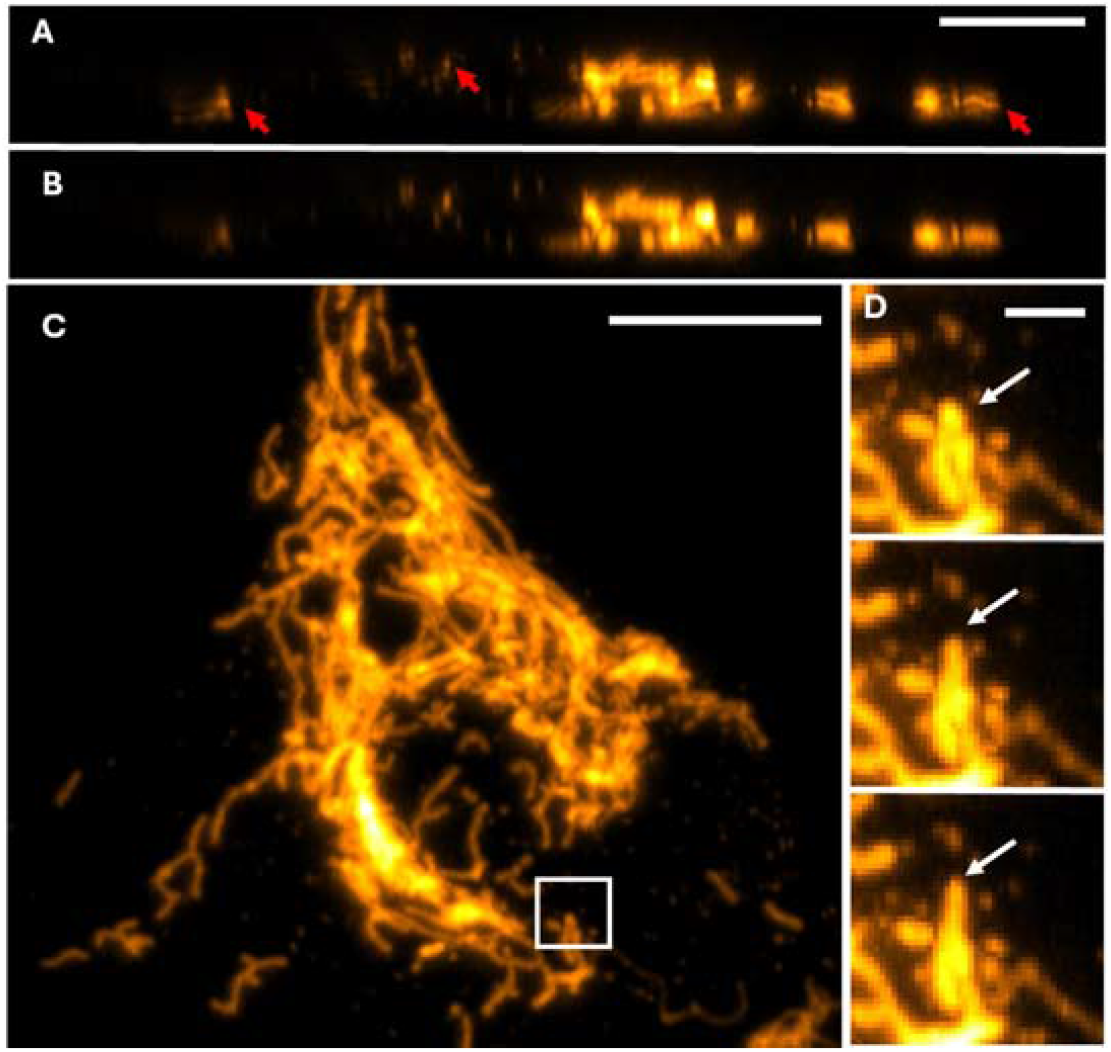
Volumetric imaging of mitochondrial dynamics at 2Hz volume rate in U2OS cells labeled with OMP25-GFP. Red arrows point at aliasing artifacts. **A** cross-sectional view of 2x under-sampled dataset. **B** Same view but using our reconstruction technique. **C** Maximum intensity projection of the U2OS cell. **D** Magnified view of the boxed region in **C**. White arrows point at an extruding mitochondrion. Scale Bar: **A, C**: 10 microns; **D**: 2 microns

**Figure 6C** shows the first timepoint of the time lapse series, and the insets in **Figure 6D** show three consecutive timepoints. The increased acquisition speed allowed us to observe rapid rearrangements of the mitochondria (**Supplementary Movie 1**).

## 3. Discussion

We have introduced a method to accelerate the volumetric acquisition rate of OPM through under-sampling and subsequent recovery of aliased information. We exploit that the major axes of the OTF are not co-aligned with the reference frame of OPM’s tertiary imaging system. As such, the tilted OTF leaves empty spaces in the rectangular Fourier domain, which can be partially filled with aliased information stemming from under-sampling. As long as there is no overlap between the primary OTF and the aliased information, we show that we can restore an OTF, and imaging performance in real space, that is close to Nyquist sampling. The degree of under sampling depends on the aspect ratio of the OTF, and as such of the PSF anisotropy. For a high-resolution system, this afforded a two-fold speed increase over critical Nyquist sampling, whereas the mesoscopic OPM, with a larger degree of anisotropy, could do fourfold faster acquisitions. Indeed, there are mesoscopic OPM systems with even higher PSF anisotropy where even higher speed gains might be possible [9]. This could further accelerate the use of OPM for whole brain and whole organism functional imaging[10].

Importantly, our reconstruction is straightforward: three (or more, see for example **Supplementary Figure S3**) copies of the Fourier transform of an under-sampled data set are concatenated in the k_z’_ direction. No sub-pixel shifts, interpolation or averaging of spectral overlap is needed.

The reconstructed OTF must be masked from residual other OTF copies. This masking step was in this work done with a top-hat function but could also involve other filter types in future work. Conceivably, the data could also be rotated into an x-y-z reference frame (of the primary objective) in real space, and axially cropped after a Fourier transform. This would in principle avoid the effect of masking and correspond simply to a down-sampling of the z-direction. We provide an alternative code to do this in our software repository.

We have introduced a heuristic stopping criterion for the under-sampling factor, namely once the axial resolution starts to get lower than the one of the ground truth. We reason this happens once the OTF copies start to touch, and one must make the filter band narrower. It is important to know that our reconstruction scheme seems to tolerate some overlap. This is because the parts of the OTF that overlap have opposite phases, which annihilate when superimposed (see **Supplementary Figure S5** and **Supplementary Note 1**). As such, the overlap leads to a shrinking of the OTF in the k_z_ direction. Thus, OPM could potentially be operated in an even faster volumetric mode at the expense of axial resolution.

In summary, we have shown that the OTF in OPM allows a certain degree of deliberate axial under-sampling, which leads to a form of aliasing that can be recovered in a loss-less and artifact free manner. In principle this should improve the spatiotemporal product of any OPM and SCAPE system while reducing the amount of raw data and sample irradiation. Our reconstruction scheme is simple and robust and can be directly integrated into the standard OPM post-processing workflow.

We hope that our scheme will accelerate the use of OPM and SCAPE systems.

## 5.1 Acknowledgment

R.F. is thankful for support from the National Institute of Biomedical Imaging and Bioengineering (grant R01EB035538) and the National Institute of General Medical Sciences (grant R35GM133522). We are thankful to the lab of Dr. Mike Henne for providing the U2OS cells.

## 5.2 Code availability

Example code to run the reconstructions shown in this manuscript are deposited here: https://github.com/AdvancedImagingUTSW/OPM-Alias

## 5.3 Data availability

Example data to run the reconstructions shown in this manuscript are deposited here: https://cloud.biohpc.swmed.edu/index.php/s/GbnjwtioTYCHDCB

## 5.4 Data acquisition

The high resolution OPM data has a p_x’_ and p_y’_ pixel size of 147nm. We acquired 3D data with 207 nm scan step size, which resulted in isotropic pixels (i.e. p_z’_=207nm*sin(45)). This resulted in Nyquist oversampling. We found that with double the step size, i.e. 414nm, we just started to see other OTF copies appear in a 3D Fourier transform of the data, but in real space, no artifacts were yet visible. We then defined this step size as the critical Nyquist sampling rate. The 2X down-sampled data was either directly acquired at 828nm step size, or we digitally down-sampled an existing stack that was acquired with 207nm step size.

The mesoscopic OPM has a p_x’_ and p_y’_ pixel size of 1.15microns, and the scanning step was set to 1.6 microns to achieve critical Nyquist sampling.

## Supplementary Information

**Figure S1.**
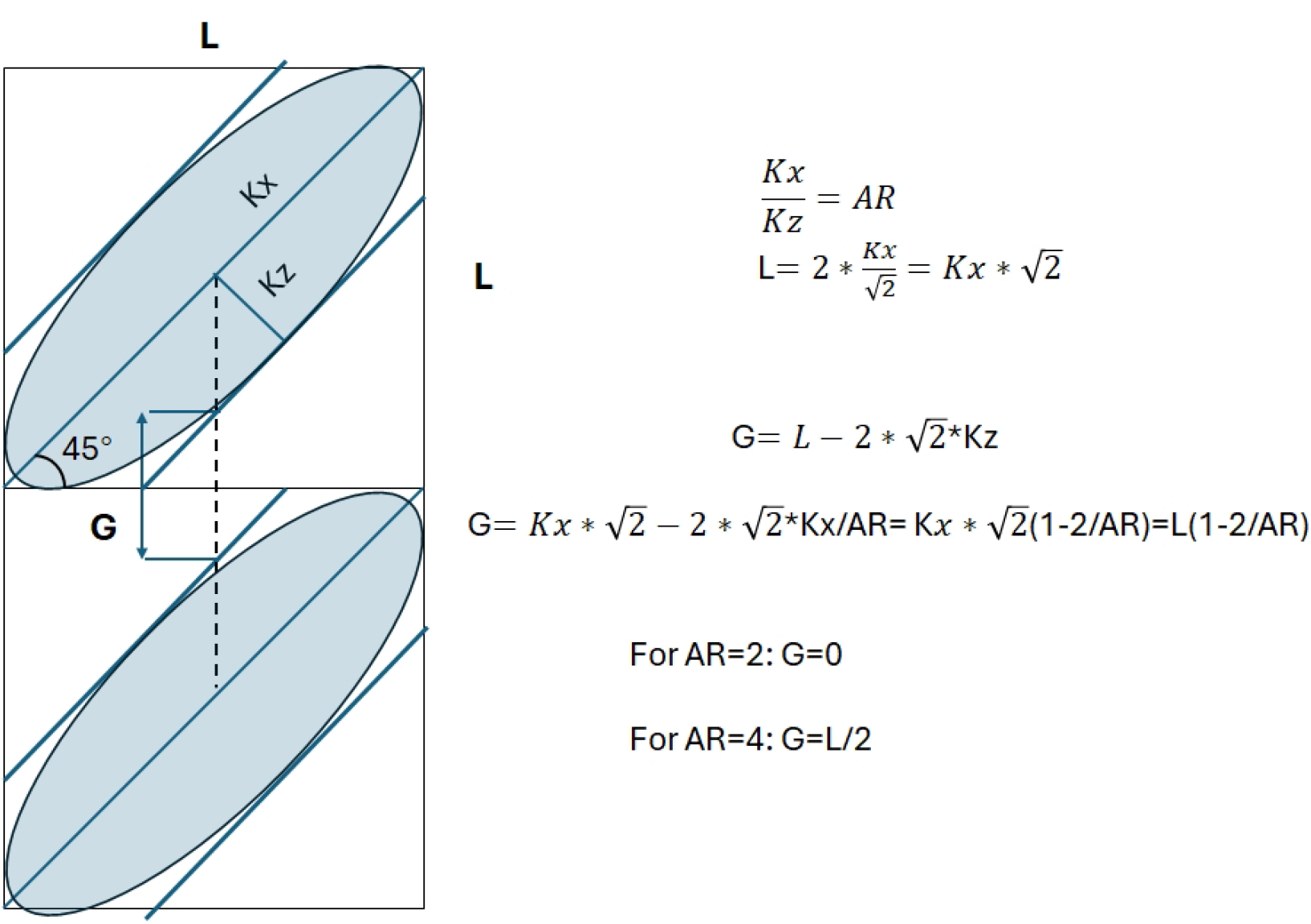
Estimation of how large the gap G between two OTF (light blue) copies is, for critical sampling and a tilt of 45 degrees. Kx and Kz are the cut-off frequencies of the OTF, and AR is the aspect ratio of the OTF.

**Figure S2.**
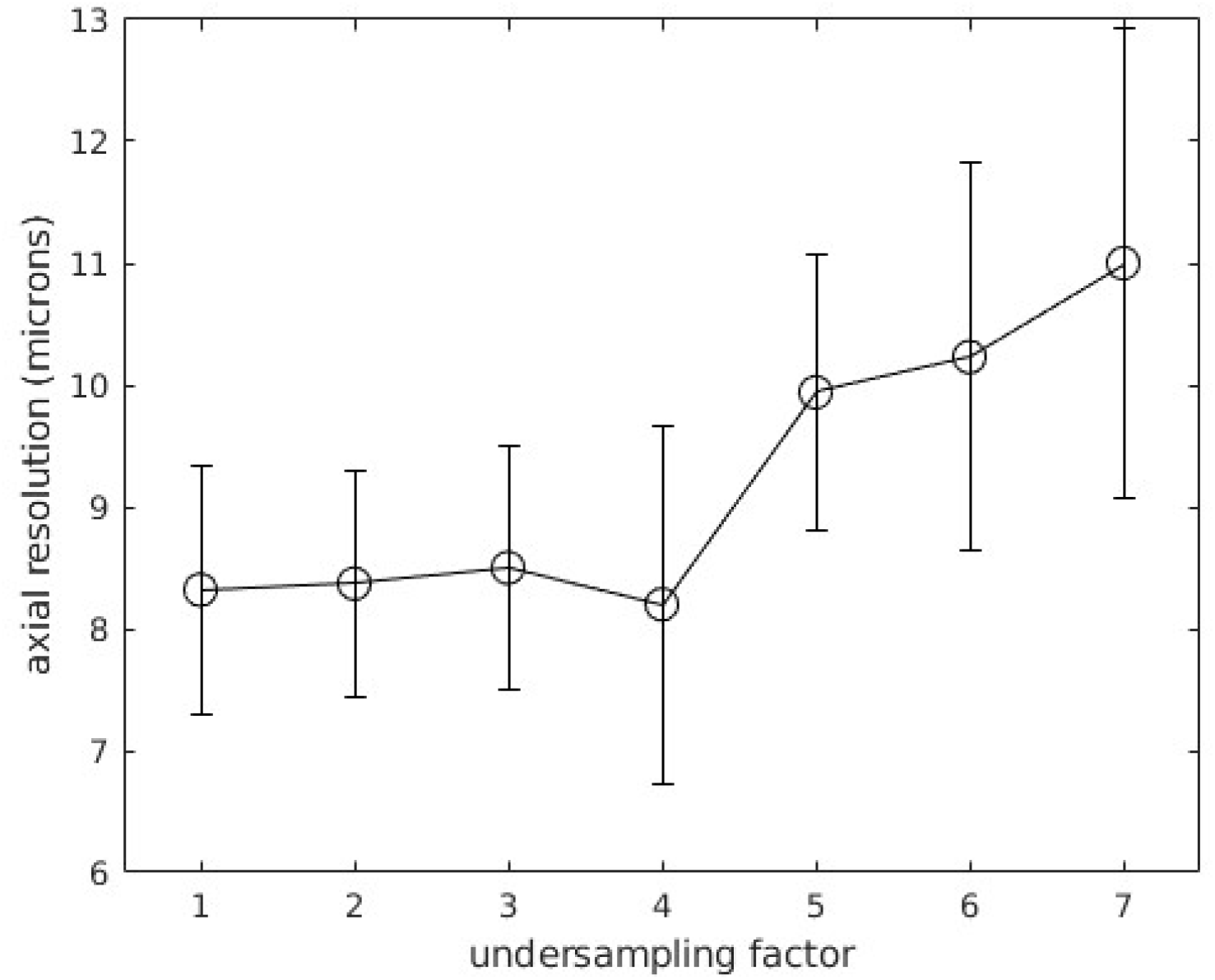
Axial resolution of the meso OPM for different under-sampling factors for the same volume of 500nm fluorescent nanospheres in Agarose. Mean values and standard deviation are shown. 1800-2000 beads were analyzed per under-sampling factor.

**Figure S3.**
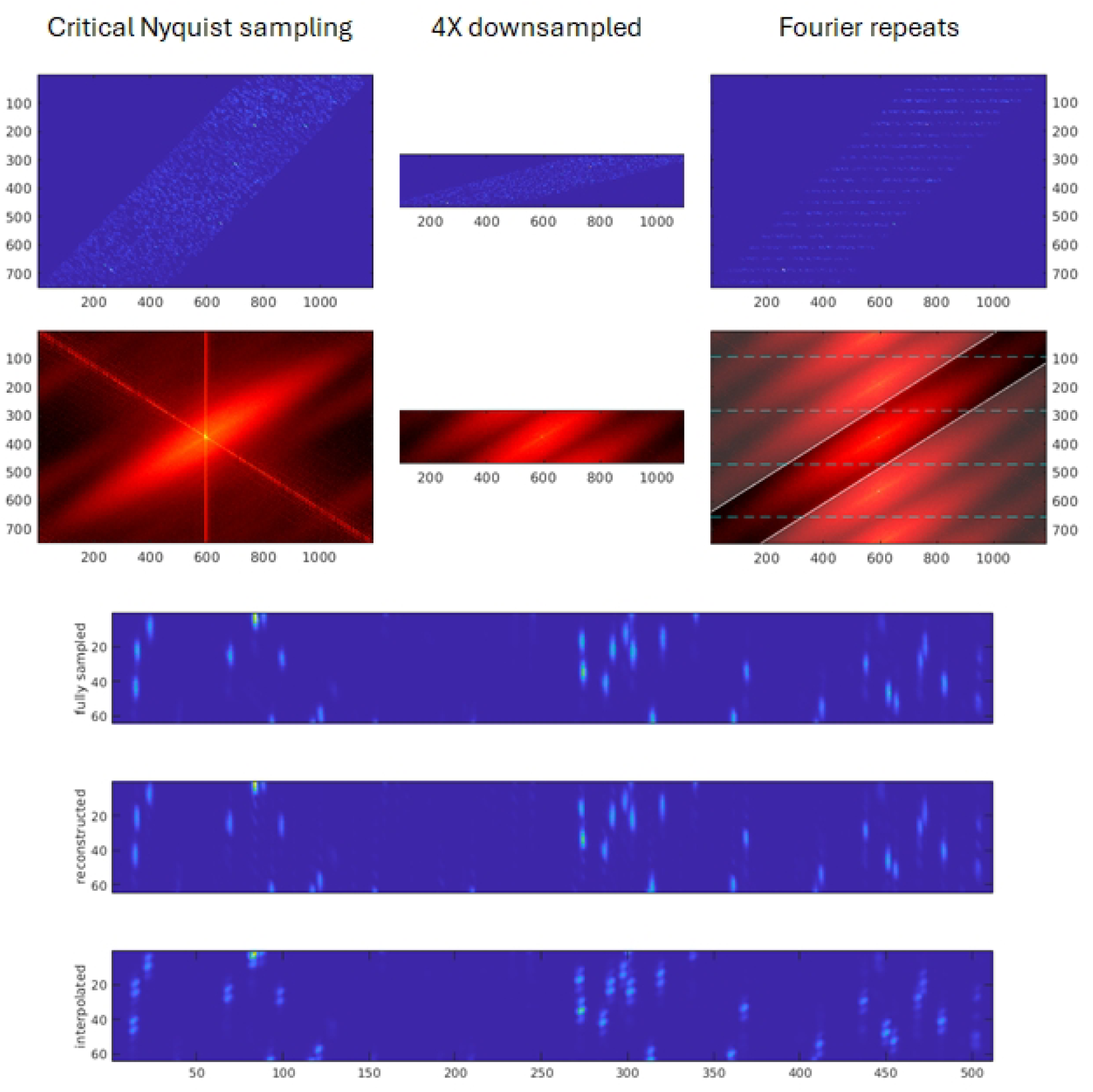
De-skewed stacks from Mesoscopic OPM imaging of 500nm fluorescent nanospheres in Agarose under critical Nyquist sampling (left), four-fold under-sampling (center), and reconstruction from under-sampled data. (right). Parula-colormap: max projections along the y-direction. Hot-colormap: Fourier transforms, log10 of the absolute value max-projected along k_y_. Reconstruction is achieved via repetition in the k_z_ direction (demarcated by blue dashed lines), followed by a hard-cut binary mask following the central OTF copy (white mask: opaque areas are zeroed out). Below shows cropped areas after rotation for the Nyquist sampled, reconstructed and interpolated cases.

**Supplementary Figure S4:**
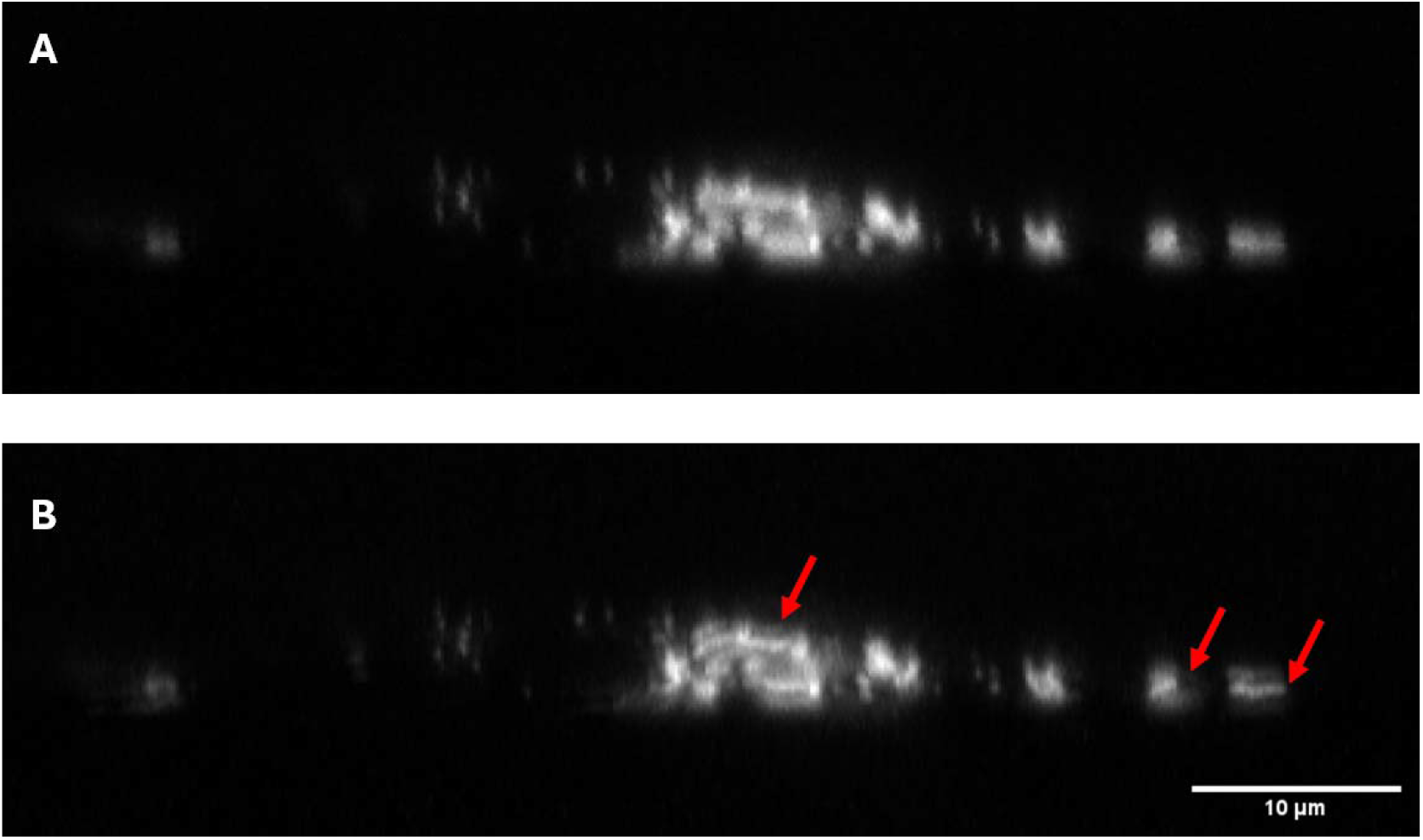
Nyquist and under-sampled mitochondria imaging in an U2OS cell. **A**: Cross sectional view of mitochondria in an U2OS cell, acquired with Nyquist oversampling. **B** Artificially down-sampling of the same dataset, and corresponding interpolated view. Arrows point at aliasing artifacts.

**Supplementary Figure S5:**
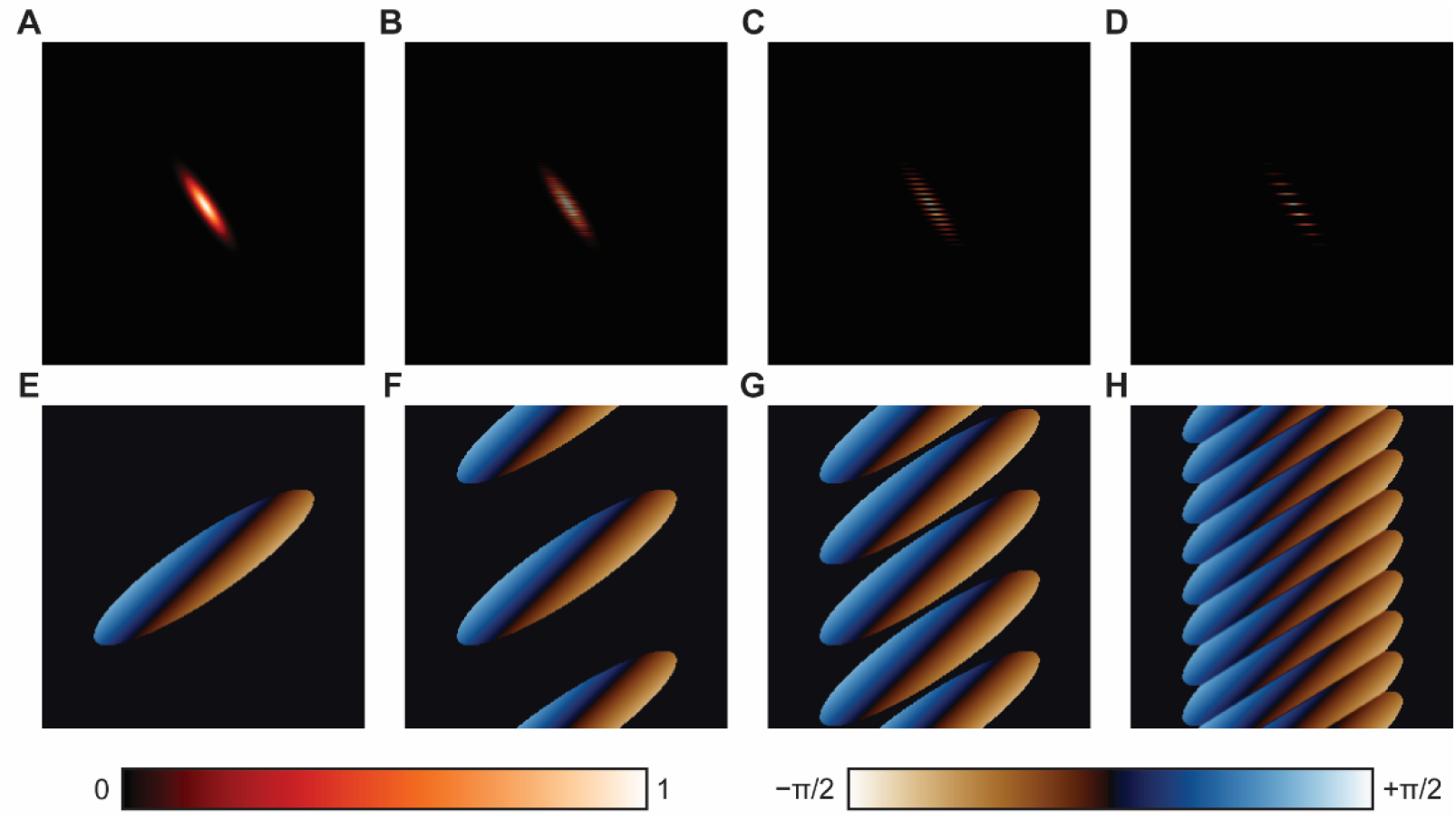
OTF annihilation via aliased copy overlap. **A**: Simulated PSF acquired with Nyquist sampling. **B** Simulated PSF with 2-fold under-sampling. **C** Simulated PSF with 4-fold under-sampling. **D** Simulated PSF with 8-fold under-sampling. **E** Phase of OTF corresponding to PSF in A. As the PSF is deliberately off-centered, a small phase ramp across the OTF is present, with negative values of phase present on the bottom right and positive values on the top left. **F** Phase of OTF corresponding to PSF in B. **G** Phase of OTF corresponding to PSF in C. **H** Phase of OTF corresponding to PSF in D. Note how each OTF is now more rectangular, as overlapping parts annihilate.

### Supplementary Note 1

To investigate the behavior of severe under sampling, PSFs were simulated as 2D Gaussian functions with principal axes at 45 degrees to the horizontal and vertical, on a 256 by 256 pixel grid (see Supplementary Figure S6). The Gaussian sigmas were 16 and 4, with the center of the function being at (130, 130). This was chosen to give a slight phase ramp across the OTF to help with visualization. For under sampled PSFs, appropriate horizontal slices within the original PSF were set to all zero. To aid visualization, the phases of the OTF were multiplied by a mask that was zero when the absolute value of the OTF was below 1E-15 to reduce spurious numerical noise. Note that in the 8-fold under sampling case, parts of the OTF copies that overlap have opposite phases and so when they add together, they annihilate. This leads to a straightforward reduction in the support of the OTF, and hence axial resolution, without introducing any artefacts after appropriate cropping of the Fourier spectrum.

## References

1. E. H. Stelzer, et al., “Light sheet fluorescence microscopy,” Nature Reviews Methods Primers 1, 73 (2021).

2. C. Dunsby, “Optically sectioned imaging by oblique plane microscopy,” Optics express 16, 20306–20316 (2008).

3. M. B. Bouchard, et al., “Swept confocally-aligned planar excitation (SCAPE) microscopy for high-speed volumetric imaging of behaving organisms,” Nature photonics 9, 113–119 (2015).

4. V. Voleti, et al., “Real-time volumetric microscopy of in vivo dynamics and large-scale samples with SCAPE 2.0,” Nature methods 16, 1054–1062 (2019).

5. M. Kumar, et al., “Integrated one-and two-photon scanned oblique plane illumination (SOPi) microscopy for rapid volumetric imaging,” Optics express 26, 13027–13041 (2018).

6. E. J. Botcherby, et al., “An optical technique for remote focusing in microscopy,” Optics Communications 281, 880–887 (2008).

7. M. G. Gustafsson, “Surpassing the lateral resolution limit by a factor of two using structured illumination microscopy,” Journal of microscopy 198, 82–87 (2000).

8. X. Ruan, et al., “Image processing tools for petabyte-scale light sheet microscopy data,” Nature Methods, 1–11 (2024).

9. M. Hoffmann and B. Judkewitz, “Diffractive oblique plane microscopy,” Optica 6, 1166–1170 (2019).

10. M. Hoffmann, et al., “Blazed oblique plane microscopy reveals scale-invariant inference of brain-wide population activity,” Nature Communications 14, 8019 (2023).

